# The *MS4A* gene cluster is a key regulator of soluble TREM2 and Alzheimer disease risk

**DOI:** 10.1101/352179

**Authors:** Yuetiva Deming, Fabia Filipello, Francesca Cignarella, Claudia Cantoni, Simon Hsu, Robert Mikesell, Zeran Li, Jorge L Del-Aguila, Umber Dube, Fabiana Geraldo Farias, Joseph Bradley, Bruno Benitez, John Budde, Laura Ibanez, Maria Victoria Fernandez, Alzheimer’s Disease Neuroimaging Initiative (ADNI), Dominantly Inherited Alzheimer Network (DIAN), Kaj Blennow, Henrik Zetterberg, Amanda Heslegrave, Per M Johansson, Johan Svensson, Bengt Nellgård, Alberto Lleo, Daniel Alcolea, Jordi Clarimon, Lorena Rami, José Luis Molinuevo, Marc Suarez-Calvet, Estrella Morenas-Rodríguez, Gernot Kleinberger, Michael Ewers, Oscar Harari, Christian Haass, Thomas J Brett, Celeste M. Karch, Laura Piccio, Carlos Cruchaga

**Affiliations:** Department of Psychiatry, Washington University School of Medicine, 660 S. Euclid Ave. B8134, St. Louis, MO 63110, USA.; Department of Neurology, Washington University School of Medicine, St Louis, MO, USA; Department of Psychiatry and Neurochemistry, Institute of Neuroscience and Physiology, the Sahlgrenska Academy at the University of Gothenburg, Mölndal, Sweden.; Clinical Neurochemistry Laboratory, Department of Neuroscience and Physiology, University of Gothenburg, Sahlgrenska University Hospital, Mölndal, Sweden.; Department of Neurodegenerative Disease, UCL Institute of Neurology, Queen Square, London, UK.; UK Dementia Research Institute at UCL, London, UK.; Department of Anesthesiology and Intensive Care Medicine, Sahlgrenska University. Hospital, Mölndal, Sweden and Institute of Clinical Sciences, the Sahlgrenska Academy at the University of Gothenburg, Sweden; Department of Internal Medicine, Institute of Medicine, the Sahlgrenska Academy at the University of Gothenburg, Göteborg, Sweden.; Neurology Dept. Hospital de la Santa Creu i Sant Pau. Sant Antoni M^a^ Claret 167. Barcelona 08025. Spain; IDIBAPS. Alzheimer’s disease and other cognitive disorders unit, Neurology Service, ICN Hospital Clinic.; Alzheimer’s disease and other cognitive disorders unit, Neurology Service, ICN Hospital Clinic i Universitari; BarcelonaBeta Brain Research Center, Pasqual Maragall Foundation. Barcelona, Spain; Biomedical Center (BMC), Biochemistry, Ludwig Maximilians-Universitat Munchen, Munich, Germany.; Munich Cluster for Systems Neurology (SyNergy), Munich, Germany; Institute for Stroke and Dementia Research, University Hospital, LMU Munich, Germany; German Center for Neurodegenerative Diseases (DZNE), Munich, Germany; Division of Pulmonary and Critical Care Medicine, Department of Internal Medicine, Washington University School of Medicine, St. Louis, United States; Hope Center for Neurological Disorders. Washington University School of Medicine, 660 S. Euclid Ave. B8111, St. Louis, MO 63110, USA.

## Abstract

Soluble triggering receptor expressed on myeloid cells 2 (sTREM2) levels in the cerebrospinal fluid (CSF) have been associated with Alzheimer disease (AD) status. TREM2 plays a critical role in microglial activation, survival, and phagocytosis; however, the pathophysiological role of sTREM2 in AD is not well understood. Understanding the role of sTREM2 in AD may help reveal biological mechanisms underlying AD and identify novel therapeutic targets. We performed a genome-wide association study (GWAS) to identify genetic modifiers of CSF sTREM2 levels. Common variants in the membrane-spanning 4-domains subfamily A (*MS4A*) gene region were associated with higher CSF sTREM2 levels (rs1582763; *P* = 1.15×10^−15^) and replicated in independent datasets. The variants associated with increased levels of sTREM2 are also associated with reduced AD risk and delayed age-at-onset. Rs1582763 influences expression of *MS4A4A* and *MS4A6A* in multiple tissues, suggesting that one or both of these genes are important for regulating sTREM2. *MS4A* genes encode transmembrane proteins that may play a role in intracellular protein trafficking in microglia. We used human macrophages to begin to test the relationship between MS4A4A and TREM2 and found that they co-localize intracellularly and that antibody-mediated targeting of MS4A4A reduces sTREM2. Thus, genetic, molecular, and cellular findings suggest that MS4A4A regulates sTREM2. These findings also provide a mechanistic explanation of the original GWAS signal in the *MS4A* locus for AD risk and indicate that TREM2 is involved in sporadic AD risk in general, not only in *TREM2* risk-variant carriers.

## Introduction

In 2013, two groups independently identified a rare variant in *TREM2* (p.R47H) that increased risk for AD almost three-fold, which is the strongest genetic risk factor for late-onset AD since the identification of *APOE ε4* 30 years earlier (*1, 2*). *TREM2* p.R47H has also been associated with clinical, imaging, and neuropathological AD phenotypes, including advanced behavioral symptoms, gray matter atrophy, and Braak staging (*3, 4*). Since then, additional rare variants in *TREM2* have been associated with AD risk including p.R62H, p.H157Y, p.R98W, p.D87N, p.T66M, p.Y38C, and p.Q33X (*1, 5–7*).

*TREM2* encodes a protein that is part of a transmembrane receptor-signaling complex essential for the immune response of myeloid cells such as microglia. TREM2 is highly and specifically expressed in microglia in the central nervous system (CNS) where it plays a key role in microglial activation, survival, and phagocytosis (*8–11*). Recent evidence from animal models of AD suggest that TREM2 has a complex relationship with the key proteins involved in AD pathogenesis, amyloid-β and tau. There is conflicting studies about the role of TREM2 in amyloid-beta (Aβ) pathology. TREM2 has been associated with phagocytosis in the presence of Aβ (*12*). Plaque-associated neuritic dystrophy was more severe (*13, 14*) and neuron loss was exaggerated in mice deficient for TREM2, in a gene-dose dependent manner (*9*). Elevating levels of TREM2 through the introduction of a human TREM2 transgene reduced pathology and rescued cognitive function in amyloid-bearing mice (*15*). On the other hand, a common phenotype across mouse models of amyloidosis is that reduction or loss of *Trem2* leads to fewer amyloid plaque-associated microglia (*9, 14, 16*). Other studies indicate that *Trem2* deficiency protects against neurodegeneration through a dampened microglial-response to tau pathology (*17, 18*).

In addition to the full-length TREM2 that makes up part of the transmembrane receptor, there is a soluble form of TREM2 (sTREM2) that may be produced by alternative splicing or proteolytic cleavage of the full-length protein (*6, 12, 19*). Three distinct transcripts for *TREM2* were found in human brain tissue, including a short transcript alternatively spliced to exclude exon 4 thus encoding the transmembrane domain predicted for sTREM2 (*6*). Recently a cleavage site was identified on TREM2 that can be targeted by ADAM10, ADAM17, gamma-secretase, and possibly other proteases, to generate sTREM2 (*20, 21*). sTREM2 is released into the cerebrospinal fluid (CSF) where it can be quantified (*12, 22–27*). CSF sTREM2 levels are hypothesized to increase in response to microglia activation due to neurodegenerative processes (*23, 26*). Our group and others have demonstrated that CSF sTREM2 levels are elevated in AD (*22, 23, 25*). Changes in CSF sTREM2 appear to occur after amyloid accumulation, beginning approximately five years before clinical symptom onset in autosomal dominant forms of AD (*24*). Additionally, CSF sTREM2 levels are positively correlated with CSF tau and ptau levels, but not with CSF Aβ_42_, suggesting that sTREM2 may be associated with pathological processes occurring after the accumulation of Aβ (*22, 23, 25*). Thus, CSF sTREM2 levels represent an important and dynamic biomarker of disease processes throughout AD pathogenesis.

Our group and others have shown that using CSF biomarkers of complex diseases as quantitative endophenotypes can identify genes that contribute to key biological pathways involved in disease (*28–30*). In this study, we aimed to identify genetic modifiers of CSF sTREM2 levels. We conducted a genome wide association study (GWAS) of CSF sTREM2 levels from more than 1,390 individuals and have successfully detected a locus within the *MS4A* gene region (11 q12.2) that shows novel genome-wide significant association with sTREM2 levels. We have also used bioinformatic, transcriptomic, and cellular approaches to test association between *MS4A* genes and TREM2. Based on these findings, we propose that the *MS4A* gene family contains a key regulator of sTREM2.

## Results

The goal of this study was to identify genetic modifiers of CSF sTREM2 levels and to better understand the role of sTREM2 in biological pathways relevant to AD. To accomplish this goal, we analyzed genetic data and CSF sTREM2 levels from 813 individuals from the Alzheimer’s Disease Neuroimaging Initiative (ADNI) in a genome-wide association study. We then sought to replicate our findings in independent datasets (n= 580).

### CSF sTREM2 levels correlates with age, sex, and AD blomarkers

We analyzed the largest dataset to date for CSF sTREM2 levels. The discovery cohort was obtained from the Alzheimer’s Disease Neuroimaging Initiative (ADNI) study (*N* = 813). The levels of CSF sTREM2 in our analyses approximated a normal distribution without transformation. There were 606 AD cases and 207 cognitively normal controls in based on clinical diagnosis, none of whom carried previously identified risk mutations in *TREM2* (Table 1).

**Table 1.**
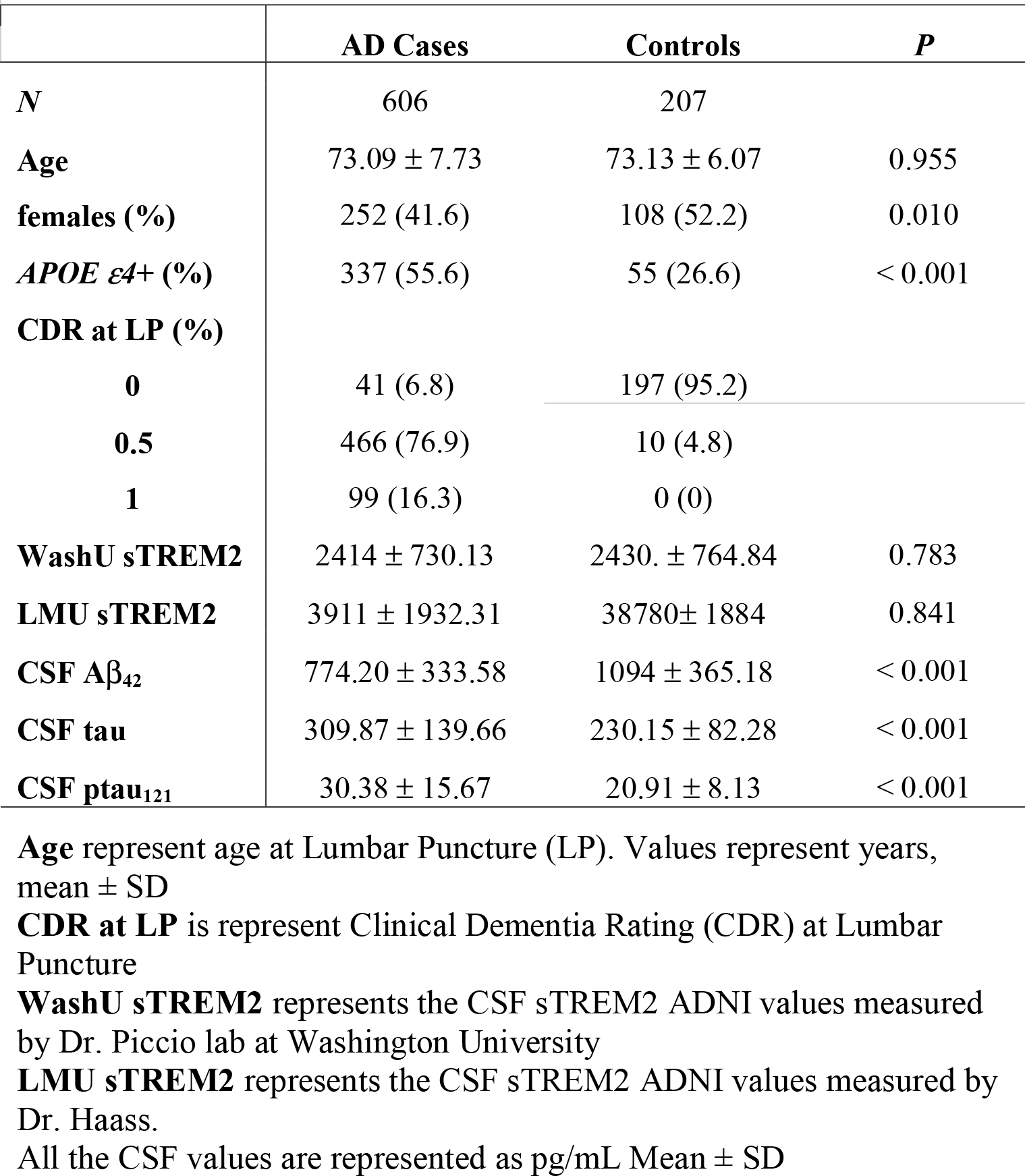
ADNI CSF cohort characteristics.

We tested the correlation between age at the time of lumbar puncture (LP) and CSF sTREM2 levels and found a positive correlation, consistent with our previous reports (Pearson *r* = 0.270, *P* = 4.66×10^−15^; Supplementary Figure S1) (*23*). We also tested CSF sTREM2 levels stratified by sex to determine whether sTREM2 levels were higher in males than females, as previously reported (*23–25, 31*). There was no significant difference between CSF sTREM2 levels in males (3,926 ± 1,969 pg/mL) versus females (3,873 ± 1,856 pg/mL, one-tailed *P* = 0.347; Fig S1). CSF sTREM2 levels were positively correlated with CSF tau (*r* = 0.377, *P* = 9.59×10^−29^) and ptau levels (r = 0.348, *P* = 2.08×10^−24^; Fig S1). Consistent with our previous findings, there was no significant correlation between CSF sTREM2 levels and CSF Aβ_42_ levels (*r* = 0.074, *P* = 0.052; Fig S1).

We also had CSF sTREM2 levels from 38 individuals heterozygous for *TREM2* risk variants which were not included in the GWAS: p.D87N, p.L211P, p.R47H, p.R62H, and p.H157Y (Table 2). There was a significant difference in CSF sTREM2 levels between *TREM2* risk variant carriers (*P* = 0.042) as reported previously (*23*). CSF sTREM2 levels were significantly lower in p.L211P carriers (2,578 ± 1,444 pg/mL) compared to non-carriers (3,903 ± 1,919 pg/mL, *P* = 0.026; Supplementary Figure S2). CSF sTREM2 levels were significantly elevated in p.R47H carriers (4,852 ± 1,172 pg/mL) compared to p.L211P carriers (*P* = 0.029) and p.D87N carriers (2,082 ± 239 pg/mL, *P* = 0.041), but they were not significantly different from non-carriers (*P* = 0.223; Fig S2).

**Table 2.** ADNI CS1f cohort characteristics by *TREM2* mutation carriers.

|  | **p.R62H** | **p.R47H** | **p.L211P** | **p.D87N** | **p.H157Y** | **P** |
| --- | --- | --- | --- | --- | --- | --- |
| *N* | 18 | 4 | 11 | 4 | 1 |  |
| **Age** | 75.01 ± 6.48 | 73.24 ± 14.51 | 72.78 ± 4.36 | 71.14 ± 5.95 | 73.10 ± NA | 0.843 |
| **Females (%)** | 9 (50.0) | 2 (50.0) | 6 (54.5) | 0 (0) | 1 (100.0) | 0.393 |
| **APOE ε4+ (%)** | 7 (38.9) | 3 (75.0) | 2 (18.2) | 2 (50.0) | 0 (0) | 0.194 |
| **CDR at LP (%)** |  |  |  |  |  | 0.815 |
| **0** | 4 (22.2) | 1 (25.0) | 5 (45.5) | 1 (25.0) | 1 (100.0) |  |
| **0.5** | 10 (55.6) | 3 (75.0) | 5 (45.5) | 3 (75.0) | - |  |
| **1** | 4 (22.2) | - | - | - | - |  |
| **2** | - | - | 1 (9.1) | 4 (0.7) | - |  |
| **Case status** |  |  |  |  |  | 0.871 |
| **AD** | 13 (72.2) | 2 (50.0) | 6 (54.5) | 1 (25.0) | - |  |
| **CO** | 3 (16.7) | 1 (25.0) | 3 (27.3) | 1 (25.0) | 1 (100.0) |  |
| **WashU sTREM2** | 2153 ± 842.34 | 2573 ± 237.61 | 1636 ± 563.94 | 1570 ± 162.55 | 3658.23 ± NA | 0.140 |
| **LMU sTREM2** | 3335 ± 1882 | 4791 ± 965.51 | 2386 ± 1389 | 1615 ± 757.86 | 5641.71 ± NA | 0.055 |
| **CSF Aβ<sub>42</sub>** | 976.76 ± 381.52 | 828.13 ± 351.90 | 1076 ± 507.09 | 685.67 ± 251.93 | NA | 0.443 |
| **CSF tau** | 303.46 ± 135.43 | 307.85 ± 97.93 | 246.24 ± 113.22 | 252.97 ± 122.51 | 213.70 ± NA | 0.678 |
| **CSF ptau<sub>121</sub>** | 29.31 ± 14.86 | 30.80 ± 13.93 | 22.18 ± 12.66 | 23.42 ± 12.99 | 18.05 ± NA | 0.729 |

**P:** is the p-value for an ANOVA analysis comparing all the variants
**Age** represent age at Lumbar Puncture (LP). Values represent years, mean ± SD
**CDR at LP** is represent Clinical Dementia Rating (CDR) at Lumbar Puncture
**WashU sTREM2** represents the CSF sTREM2 ADNI values measured by Dr. Piccio lab at Washington University
**LMU sTREM2** represents the CSF sTREM2 ADNI values measured by Dr. Haass.
All the CSF values are represented as pg/mL Mean ± SD

### The MS4A gene region is associated with CSF sTREM2 levels

To identify genetic variants that influence CSF sTREM2, we tested for association between CSF sTREM2 levels and single-nucleotide polymorphisms (SNPs) with minor allele frequency (MAF) >0.02 using an additive linear regression model with age, sex, and two principal components (PCs) as covariates. A total of 7,320,475 genotyped and imputed variants passed strict quality control as described in the methods. *TREM2* risk-variant carriers were excluded from the GWAS analyses.

A genome-wide significant genetic association with CSF sTREM2 levels was identified on chromosome 11 within the *MS4A* gene region (Figure 1 and Supplementary Figure S3). One of the top SNPs was rs1582763, an intergenic variant nearest *MS4A4A* (located 26 Kb 5 ◻ 11q12.2). This common variant (A allele, MAF = 0.368, genotyped) produced a genome-wide significant association with CSF sTREM2 levels (*N* = 807, *β* = 735.1, *P* = 1.15×10^−15^; Figure 1). This variant explains more than 6% of the variance in sTREM2 levels (adjusted r^2^ = 0.064; see supplementary results). Results from case-control stratified analyses indicate that both clinically diagnosed AD cases (*N* = 606, *β* = 675.8, *P* = 8.19×10^−10^) and cognitively normal controls (*N* = 207, *β* = 912.4, *P* = 5.20×10^−8^) contributed to the association between the *MS4A* locus and CSF sTREM2 levels (Supplementary Figure S4). All imputed and genotyped SNPs in LD with rs1582763 passed the suggestive significance threshold in the single-variant analysis (Supplementary Table S1, *P* < 1×10^−5^). No genome-wide or suggestive signals were found in the *APOE*, *ADAM10* or *ADAM17*regions (Supplementary Figure S5).

To determine whether there was more than one independent signal within the *MS4A* locus, we performed conditional analyses including rs1582763 in the regression model. After conditioning for rs1582763, the most significant variant was rs6591561 (G allele, MAF = 0.316, genotyped), a missense variant within *MS4A4A* (NP_076926.2:p.M159V), which is not in high LD with rs 1582763 (*r^2^* = 0.112, D◻ = 0.646). Rs6591561 (*MS4A4A* p.M159V) was associated with CSF sTREM2 levels but in the opposite direction of the main signal (rs1582763). The minor allele of rs6591561 was associated with lower CSF sTREM2 levels, and this association is independent of rs1582763 (before conditioning: *β* = −783.7, *P* = 1.47×10^−9^; after conditioning: *β* = −378.4, *P* = 1.55×10^−4^; Fig. 1). After conditioning for rs6591561 (*MS4A4A* p.M159V), rs1582763 remained genome-wide significant (*β* = 625.7, *P* = 4.52×10^−11^; Fig 1), providing further evidence that these are independent signals.

**Figure 1.**
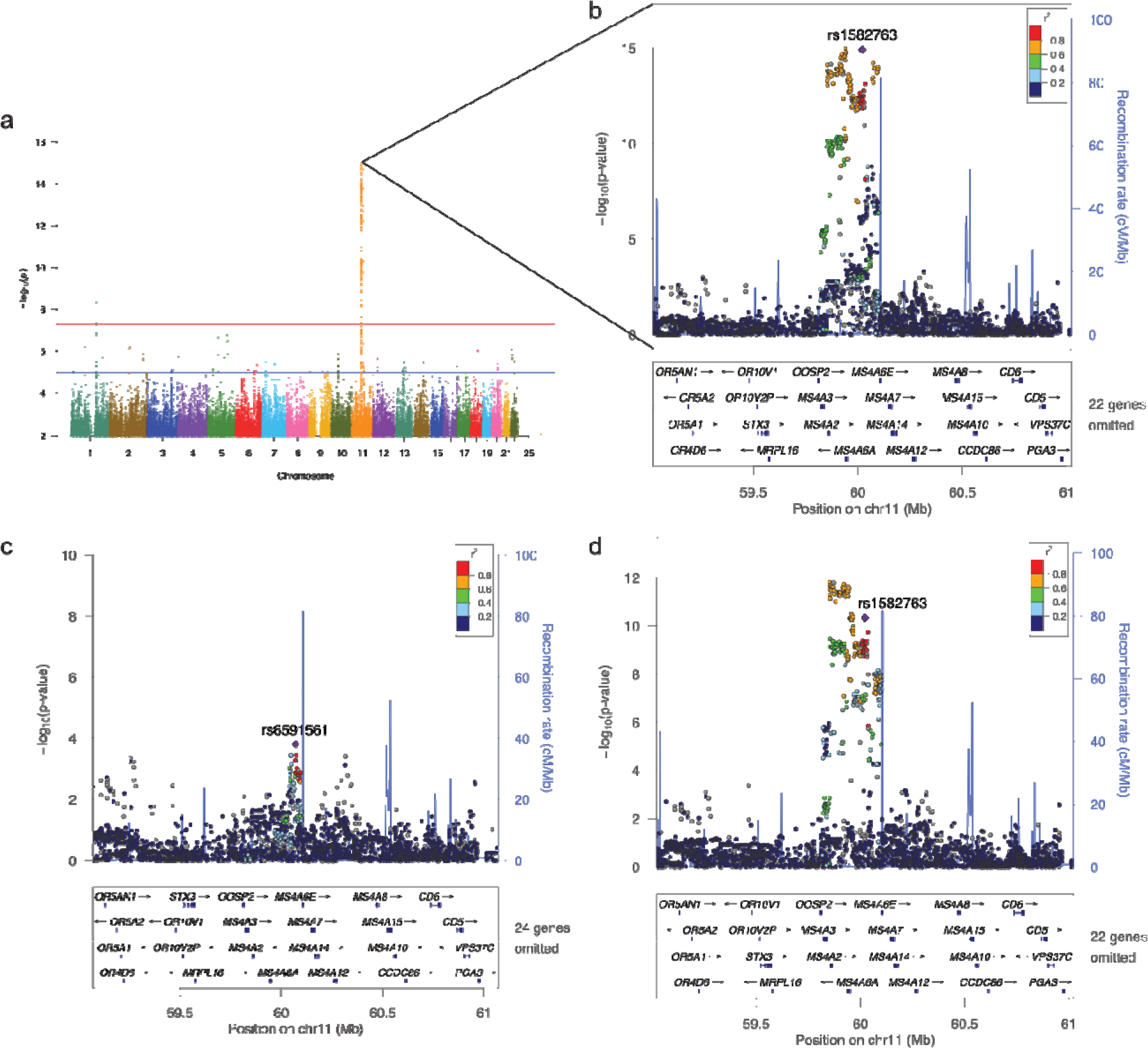
Association plots from single-variant analysis of CSF sTREM2 levels. **(a)** Manhattan plot shows the negative log_10_-transformed *P*-values on the y-axis for CSF sTREM2 levels based on the LMU measures (see material and methods). Comparable results were found with the WU measures (see supplementary material). The horizontal lines represent the genome-wide significance threshold (red) and suggestive threshold (blue). **(b)** Regional association plot of the *MS4A* gene region in the single-variant analysis. **(c)** Regional association plot of the *MS4A* region after conditioning on the top SNP (rs1582763) and **(d)** after conditioning on rs6591561 (*MS4A4A* p.M159V). The SNPs labeled on each regional plot are represented by a purple diamond. Each dot represents individual SNPs and dot colors in the regional plots represent LD with the named SNP. Blue vertical lines in the regional plots show recombination rate as marked on the right-hand y-axis.

### Replication in independent datasets and meta-analysis

To replicate the associations of rs1582763 and rs6591561(MS4A4A p.M159V), we analyzed additional 580 samples with CSF sTREM2 levels and genetic data that were obtained from six different studies (Charles F and Joanne Knight Alzheimer’s Disease Research Center, the Dominantly Inherited Alzheimer Network, two studies from the Sahlgrenska Academy at the University of Gothenburg (GHDEM and GHPH), the Memory Unit and Alzheimer’s laboratory at the Hospital of Sant Pau in Barcelona (Sant Pau Initiative on Neurodegeneration; SPIN), and the ICN Hospital Clinic-IDIBAPS in Barcelona. A description of these datasets can be found elsewhere (*23–25, 31*) and in Supplementary Table S2.

We replicated the association of rs1582763 with elevated CSF sTREM2 levels (*z* = 5.743, *P* = 9.28×10^−9^; Figure 2). A meta-analysis including the ADNI and the replication dataset showed a strong association between rs1582763 and CSF sTREM2 levels (*P*_*meta*_ = 4.48×10^−21^; Fig 2). Additionally, we replicated the association between rs6591561 (*MS4A4A* p.M159V) and reduced CSF sTREM2 levels in the independent dataset (*z* = −3.715, *P* = 2.03×10^−4^). A meta-analysis with the ADNI discovery dataset provided a p-value of *P* = 1.65×10^−u^. Individual results for each dataset are shown in Supplementary Table S3.

**Figure 2.**
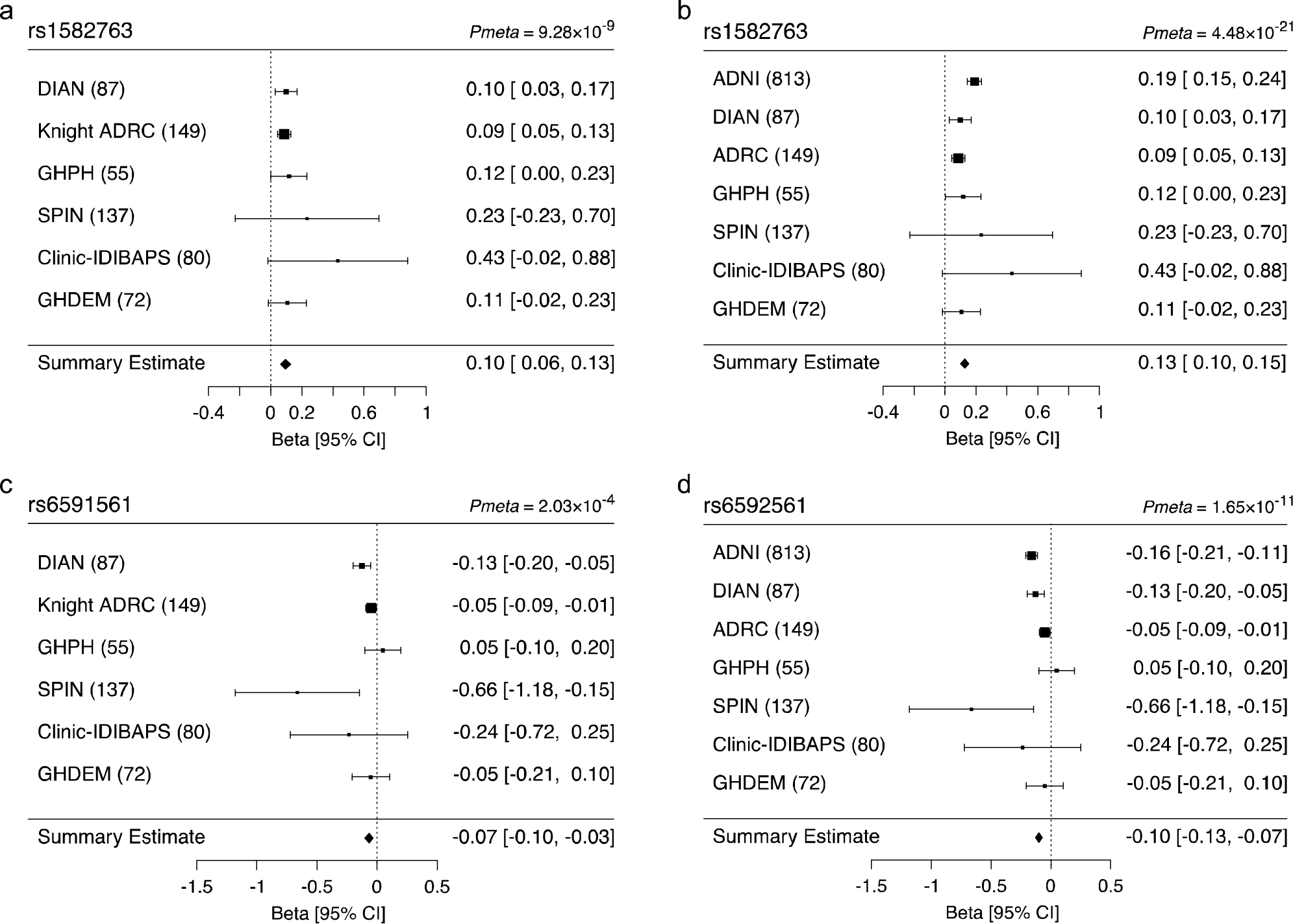
Meta-analysis of replication data sets. **(a)** Forest plot of the meta-analysis of rs1582763 in the replication datasets from DIAN, Knight ADRC, GHPH, SPIN, Clinic-IDIBAPS, and GHDEM; **(b)** forest plot of the metaanalysis of rs1582763 including ADNI; **(c)** forest plot of p.M159V (rs6591561) in the meta-analysis of DIAN, Knight ADRC, GHPH, SPIN, Clinic-IDIBAPS, and GHDEM; and **(d)** including ADNI.

### Comparisons between MS4A association and APOE

*APOE* genotype is the strongest genetic risk factor for AD (*32*) and *APOE ε4* is the strongest genetic association for CSF Aβ_42_ and tau levels (*30, 33*). To determine whether *APOE* genotype is associated with CSF sTREM2 levels, we created a variable representing *APOE*-mediated AD risk by recoding *APOE* genotype for each individual as 0 for *ε2/ε2*, 1 for *ε2/ε3*, 2 for *ε3/ε3*, 3 for *ε2/ε4*, 4 for *ε3/ε4*, and 5 for *ε4/ε4*. We added this *APOE* genotype risk variable to a regression model, adjusting for age, sex, and two PCs for population stratification, and found that *APOE* genotype was not significantly associated with CSF sTREM2 levels (*β* = −26.5, *P* = 0.613). The SNP commonly used as a proxy for *APOE ε4*, rs769449, was also not associated with CSF sTREM2 levels (*β* = −106.9, *P* = 0.308).

The influence of APOE genotype on Aβ pathology, and thereby biomarkers for Aβ pathology, including CSF Aβ42, has been a consistent finding (*30, 34*). Of the 813 individuals in this study, we had data for CSF Aβ_42_ levels available in 695 individuals. To compare CSF Aβ_42_ and sTREM2 levels on the same scale, we converted Aβ_42_ and sTREM2 values to z-scores by subtracting the mean and dividing by the standard deviation for each respective protein. After conversion, we verified the association between *APOE* genotype and CSF Aβ_42_ levels. As expected, there was a strong negative effect of *APOE* genotype on CSF Aβ_42_ levels (*β* = −0.361, *P* = 1.69×10^−34^). We also stratified the CSF Aβ_42_ levels in quartiles and calculated the OR of *APOE* genotype for the first vs the last quartile. *APOE* genotype had an OR = 2.84 for lower CSF Aβ_42_ levels (95% CI = 2.30 − 3.56, *P* = 8.44×10^−21^).

Within the same subset of individuals, the effect size of rs1582763 on CSF sTREM2 levels was similar (*β* = 0.366, *P* = 7.53×10^−13^) to that of *APOE* on CSF Aβ_42_ levels (*β* = −0.361, *P* = 1.69×10^−34^). When comparing the lower vs. upper quartiles of CSF sTREM2 levels, rs1582763 had an OR = 3.34 for higher CSF sTREM2 (95% CI = 2.37 - 4.82, *P* = 2.50×10^−11^), which is similar to the effect of *APOE* on CSF Aβ_42_ levels (OR = 2.84, 95% CI 2.30 - 3.56). Thus, the impact of rs1582763, in the *MS4A* gene region, on CSF sTREM2 levels is similar to that of *APOE*, the major regulator of AD risk, on CSF Aβ_42_ levels.

### Additional genes associated with CSF sTREM2 levels

To identify genes associated with CSF sTREM2 levels, we used MAGMA, which maps every SNP to the closest gene, takes into account the LD structure, and uses multiple regression analyses to provide a p-value for the association of each gene with CSF sTREM2 levels. There were four genes associated with CSF sTREM2 levels and passed multiple test correction (*P* < 2.75×10^−6^) All of those genes belong to the *MS4A* gene family. and included *MS4A4A* (*P* = 3.15×10^−11^; Figure 3), *MS4A4E*(*P* = 6.13×10^−12^), foll*MS4A2*(*P* = 1.29×10^−11^), and *MS4A6A* (*P* = 1.44×10^−11^). However there were also other genes, specifically, *TREML2* that was near gene-wide significance with 89 mapped variants (*P* = 3.23×10^−6^; Fig 3), suggesting that there are other genes and regions that are associated with sTREM2 levels, although the effect size is not as large as the effect size for the *MS4A* gene region

**Figure 3.**
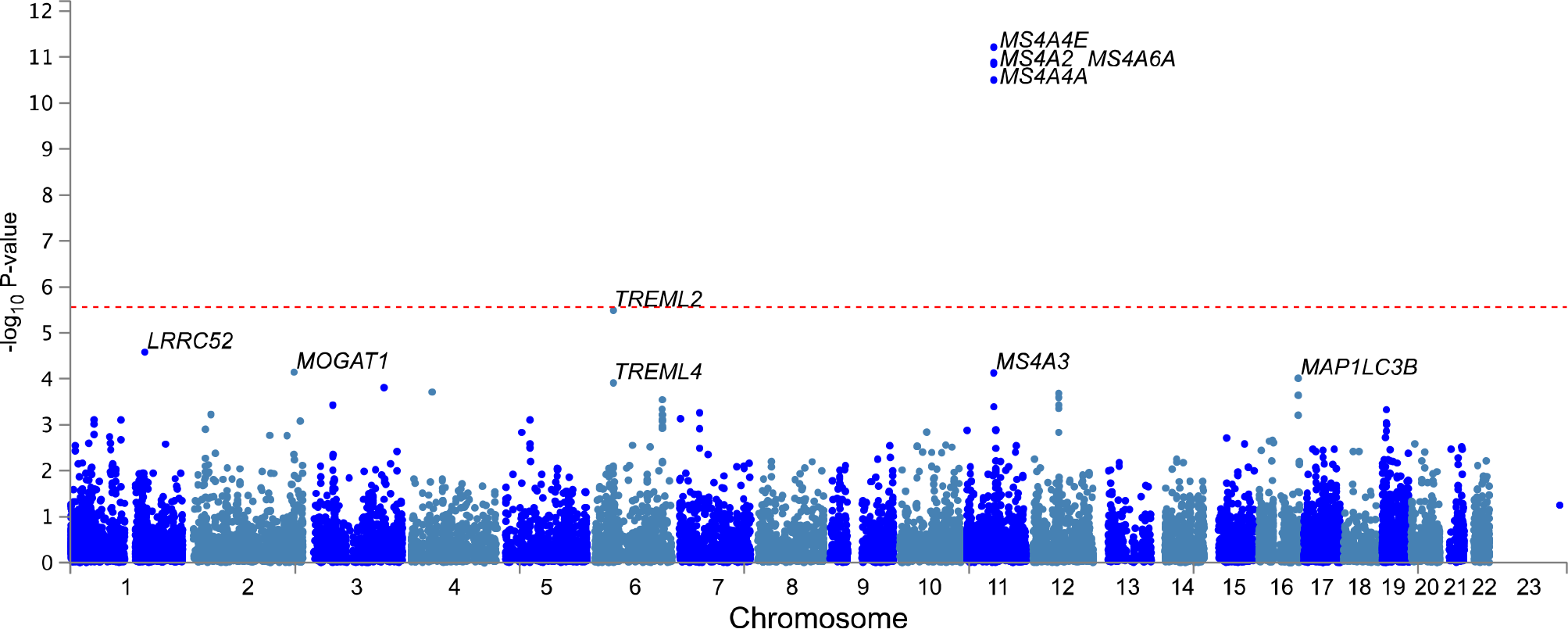
Association plot from gene-based analysis of CSF sTREM2 levels. Manhattan plot shows the negative log_10_-transformed p-values on the y-axis. Each dot represents one gene; the top ten genes in the MAGMA results are labeled. The red dotted line represents the genome-wide significance threshold (*P* = 2.74×10^−6^).

### Functional annotation of genome wide significant signals in the MS4A gene region

The genome-wide significant association with CSF sTREM2 levels is located in a gene-rich region which includes at least 15 genes, most of which are members of the *MS4A* gene family including *MS4A2*, *MS4A6A*, *MS4A4E*, *MS4A4A*, *MS4A6E*, *MS4A7*, *MS4A14*, *MS4A5*, *MS4A1*, *MS4A12*, *MS4A13*, and *MS4A8*. In order to determine the functional variants and genes that regulate CSF sTREM2 levels, we performed additional bioinformatic analyses.

We first examined whether any of the genome-wide significant variants were located within exonic regions and identified two coding variants: the missense variant within *MS4A4A* rs6591561 (p.M159V, MAF = 0.316, *β* = −593.6, *P* = 1.47×10^−9^), which is reported above, and a synonymous variant within *MS4A6A*, rs12453 (p.L137L; MAF = 0.392, *β* = 710.5, *P* = 1.77×10”^−15^) which is in LD with rs1582763 (*r^2^* = 0.782, D◻ = 0.924). Then we tested if these SNPs showed evidence of expression quantitative trait locus (eQTL) effects on *MS4A* genes. Rs6591561 has a modest *cis*-eQTL effect on both *MS4A4A* (*Z* = 3.16, *P* = 1.60×10^−3^) and *MS4A6A* (*Z* = 3.49, *P* = 4.90×10^−4^) in whole blood (*35*). The synonymous variant, rs12453, appears to exert stronger *cis*-eQTL effects on *MS4A2* (Z = 9.09, *P* = 9.98×10^−20^), *MS4A4A* (Z = −7.28, *P* = 3.46×10^−13^), and *MS4A6A* (Z = −25.80, *P* = 8.08×10^−147^) in whole blood (*35*).

We also searched for eQTL effects of rs1582763, the top SNP for CSF sTREM2 levels. Rs1582763 produced an eQTL effect on *MS4A6A* in whole blood (GTEx: *β* = −0.089, *P* = 3.90×10^−5^; Westra Blood: *Z* = −23.42, *P* = 2.95×10^−121^). Rs1582763 also influences expression of *MS4A4A* (Westra Blood: *Z* = −7.21, *P* = 5.52×10^−13^) and *MS4A2*(Westra Blood: *Z* = 8.06, *P* = 7.49×10^−16^). Using gene expression data from blood obtained from 365 individuals in the ADNI studies, we verified that rs1582763 has a *cis*-eQTL effect on *MS4A4A* (*β* = −0.226, *P* = 1.04×10^−4^) and *MS4A6A* (*β* = −0.080, *P* = 6.02×10^−6^). The eQTL effect of rs1582763 on *MS4A4A* and *MS4A6A* was observed in both clinically diagnosed AD cases (*N* = 235; *MS4A4A*: *β* = −0.194, *P* = 6.05×10^−3^; *MS4A6A*: *β* = −0.053, *P* = 1.24×10^−2^) and cognitively normal controls (*N* = 80; *MS4A4A*: *β* = −0.326, *P* = 1.10×10^−2^; *MS4A6A*: *β* = −0.138, *P* = 3.32×10^−4^). Together, this data suggests that the principle genes driving the GWAS signal for CSF sTREM2 levels may be *MS4A4A*, *MS4A6A*, or both.

### MS4A genes and TREM2 gene expression are highly correlated

We tested for correlations between expression of *TREM2* and members of the *MS4A* family using brain RNA-seq data from the Knight ADRC (*N* = 103) and DIAN (N = 19) (*36*). Among the 16 genes tested from the *MS4A* family, the expression of three genes (*MS4A4A*, *MS4A6A*, and *MS4A7*) was consistently positively correlated with *TREM2* gene expression in human brain tissue (Supplementary Figure S6 and Supplementary Table S4). In autopsy-confirmed late-onset AD cases and controls (Knight ADRC), *TREM2* expression was highly correlated with expression of *MS4A4A* (*N* = 40, *r* = 0.41, *P* = 8.00×10^−3^) and *MS4A6A* (*N* = 41, *r* = 0.67, *P* = 1.60×10^−6^; Fig S6 and Table S4). These findings were replicated in two independent RNA-seq datasets obtained from the Mayo Clinic Brain Bank (*N* = 162; *MS4A4A*: *r* = 0.68, *P* = 1.00×10^−23^; *MS4A6A*: *r* = 0.76, *P* = 1.60×10^−31^; Supplementary Figure S7 and Supplementary Table S5) and Mount Sinai Brain Bank (*N* = 300; *MS4A4A*: *r* = 0.61, *P* = 1.60×10^−16^; *MS4A6A*: *r* = 0.52, *P* = 7.90×10^−8^; Supplementary Figure S8 and Supplementary Table S6). These data further support that *MS4A4A* or *MS4A6A* may be the principle genes regulating CSF sTREM2 levels.

### Targeting MS4A4A on human macrophages decreases levels of soluble TREM2 in vitro

*MS4A4A* and *MS4A6A* are both highly expressed in microglia (*37*). However, the mouse ortholog for *MS4A6A* is *Ms4a6e* and there is no mouse ortholog for *MS4A4A* (*38, 39*). Thus, to understand the role of *MS4A* genes on TREM2 function, it is essential to use human models. To begin evaluating the functional connection between TREM2 and MS4A4A and MS4A6A, we examined human macrophages, which are myeloid cells expressing TREM2 and can serve as a surrogate for microglia (*40*).

To examine the therapeutic potential of targeting these MS4A proteins on sTREM2 levels, we used an antibody-based approach. Human macrophages were derived *in vitro* from blood monocytes and media from the cells was tested for the presence of sTREM2 by ELISA in the presence or absence of the cytokine IL-4 as previously described (*26*). IL-4-mediated macrophage stimulation results in upregulation of TREM2 and MS4A4A (*41*). sTREM2 was detected in the macrophage media beginning at day 3-post isolation and sTREM2 levels increased progressively with time in culture (Supplementary Figure S9). On day 7, stimulation with IL-4 significantly increased levels of sTREM2 in the media (Figure 4A). Next, we evaluated whether antibody-mediated targeting of MS4A4A or MS4A6A would impact sTREM2 levels. Human macrophages were treated with commercially available antibodies to MS4A4A, MS4A6A, and isotope controls (see methods for details) on day 7 and sTREM2 was measured in the media 48 hours after treatment. We found that antibodies directed to MS4A4A [N-terminal cytoplasmic domain (A) and extracellular domain (B)] were sufficient to significantly reduce extracellular sTREM2 levels (Fig. 4B). Treatment of human macrophages with an MS4A6A antibody (N-terminal cytoplasmic domain) failed to alter sTREM2 levels (Fig. 4B). We observed similar results when macrophages were stimulated with IL-4 (Fig. 4B). Thus, targeting MS4A4A with specific antibodies is sufficient to modify sTREM2 levels under basal conditions and with stimulus. However, due to limited availability of MS4A6A reagents, we cannot yet exclude the possibility that antibodies engaging extracellular epitopes on MS4A6A could also impact sTREM2 levels.

**Figure 4.**
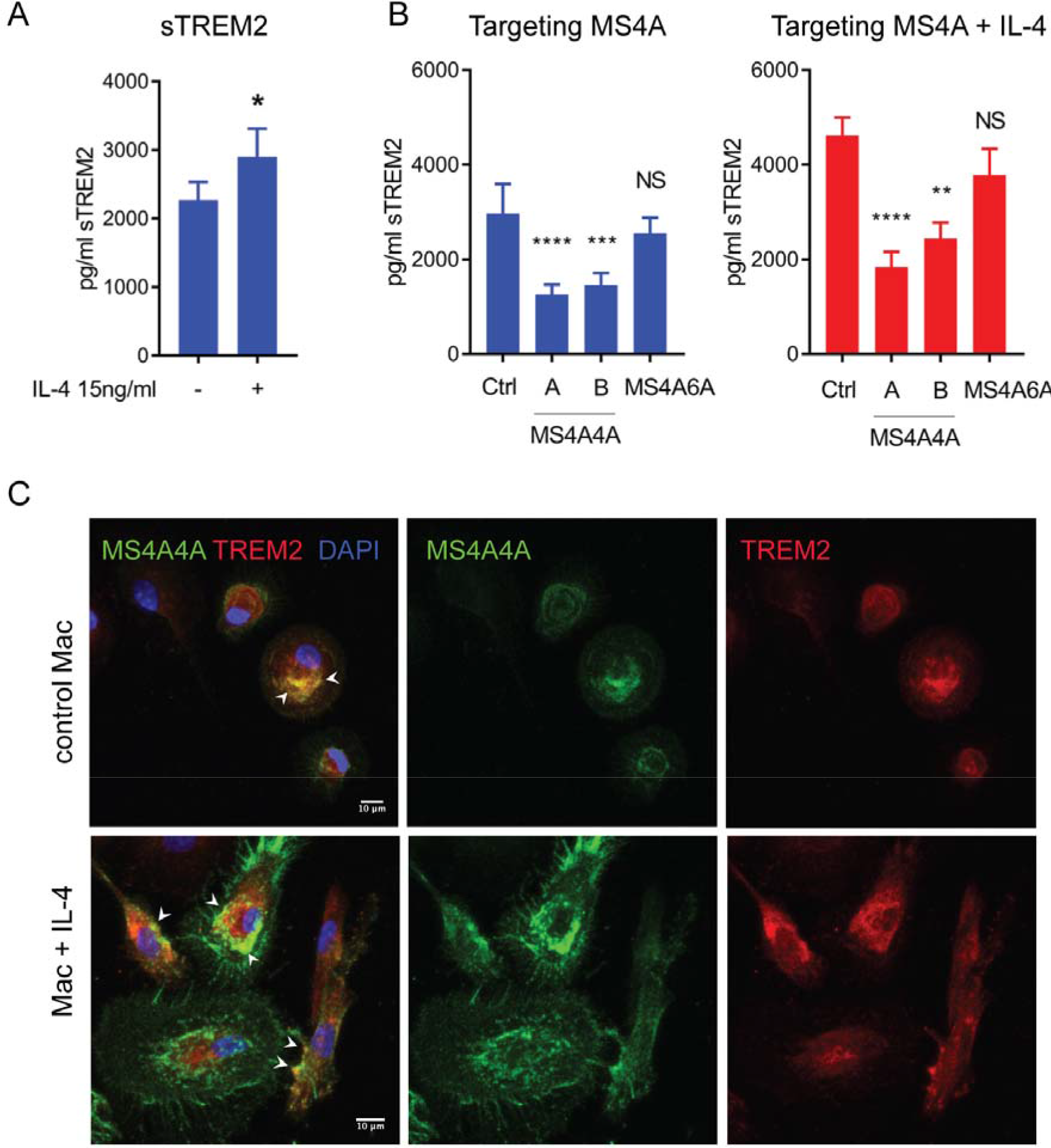
Treatment with anti-MS4A4 antibodies of human macrophages in culture leads to reduction on levels of sTREM2 in cell culture supernatants. (A) Levels of sTREM2 measured by ELISA in the cell media of human macrophages on day 7 in culture with or without stimulation with IL-4. (B) Effect on levels of sTREM2 in cell culture media by treatment with antibodies against MS4A4A and MS4A6A for 48 h. Two different anti-MS4A4 antibodies were tested, antibody A (Abcam) directed against the N-Terminal cytoplasmic domain or antibody B (Biolegend) directed against the extracellular domain of the protein. These analyses were done without or with IL-4 in culture. Bars are means ± SD. *p<0.05; **p<0.005; ***p<0.005. Pvalues were calculated by Mann-Whitney test or Kruskal Wallis H test for multigroup comparison. (C) Confocal images of human macrophages treated or not with IL-4 for 48 hours and stained for TREM2 (red) and MS4A4A (green) antibodies. Arrowheads (white) indicate colocalizing signal. Scale bar 10 μm.

### MS4A4 co-localizes with TREM2 in human macrophages

To determine whether MS4A4A are localized within similar cellular compartments with TREM2, we performed immunocytochemistry in human macrophages. MS4A4A has been previously shown to localize on monocyte/macrophages plasma membrane (*41*) and on the plasma membrane and intracellular organelles in human mast cells, where it may play a role in intracellular protein trafficking (*42*). While we observed some MS4A4A staining at the plasma membrane in the human macrophages, we detected the strongest signal in intracellular organelles mainly localized in the juxta-nuclear region (Fig. 4C). Confocal analysis revealed that MS4A4A and TREM2 co-localize in the same intracellular structures and, to a lesser extent, on the plasma membrane (Fig. 4C). Consistent with the sTREM2 findings and previous reports (*40, 41*), IL-4-mediated stimulation of the human macrophages significantly increased expression of both MS4A4A and TREM2 (Fig. S9).

## Discussion

In this study, we performed a genome-wide analysis for genetic modifiers of CSF sTREM2 levels. We observed two independent signals in the *MS4A* gene region that pass genome-wide significance thresholds for association with CSF sTREM2 levels: rs1582763 and rs6591561 (*MS4A4A* p.M159V). Importantly, rs1582763 is associated with elevated CSF sTREM2 levels and reduced AD risk (*43*) and delayed age-at-onset (*39*). Conversely, rs6591561 encoding *MS4A4A* p.M159V is associated with reduced CSF sTREM2 levels and increased AD risk and accelerated age-at-onset (*39*). These SNPs modify *MS4A4A* and *MS4A6A* in human blood and brain tissue and are highly correlated with *TREM2* gene expression. In human macrophages, MS4A4A and TREM2 co-localize in intracellular compartments and antibody-mediated targeting of MS4A4A results in a significant reduction of sTREM2. Thus, we present genetic, molecular, and functional studies that suggest that MS4A genes are key regulators of sTREM2.

Our findings that variants in the *MS4A* gene region are associated with CSF sTREM2 levels provides a biological connection between the MS4A gene cluster, TREM2, and AD risk. Prior to this study, TREM2 has been directly implicated in AD through the identification of rare variants that increase AD risk (*1, 2*). However, the role of TREM2 in sporadic disease was less clear. Common variants in the *MS4A* locus are associated with AD risk (rs1582763, OR = 0.898, *P* = 1.81×10^−15^) (*43*) and delayed age-at-onset (HR = 0.929, *P* = 9.26×10^−9^) (*39*), but the functional variant and mechanism of action has been unknown until now. Here, we show that variants within this AD risk locus modify CSF sTREM2 levels. Previous research has already suggested that changes in sTREM2 levels, or TREM2 biology in general, are likely involved in disease pathology (*12–18, 24*). These *MS4A* variants also impact *MS4A4A* and *MS4A6A* expression in human blood and brain tissue, as previously reported (*44, 45*).

The two independent signals in the *MS4A* gene region have opposing effects on CSF sTREM2 levels, which has important implications for MS4A and TREM2 biology in AD pathogenesis. Interestingly, studies in AD mouse models suggest that loss of *Trem2* has opposing effects on amyloid- and tau-mediated pathology (*14, 16, 46*). How CSF sTREM2 levels change throughout human disease remains poorly understood and is an area of active investigation. However, our findings that a SNP, rs1582763, which is associated with elevated CSF sTREM2 levels is also associated with reduced AD risk and delayed age-at-onset point to a protective role for sTREM2 in AD. Consistently, the second independent SNP, rs6591561, is associated with reduced CSF sTREM2 levels, increased AD risk, and accelerated age-at-onset, suggesting that lower CSF sTREM2 is associated with a more severe disease course. We cannot say yet whether CSF sTREM2 levels are a biomarker for inflammatory or other non-specific processes in the brain, or whether CSF sTREM2 levels more directly contribute to disease pathology.

We still have a poor understanding of the origin and function of sTREM2. Three distinct transcripts for *TREM2* were found in human brain tissue, including a short transcript alternatively spliced to exclude exon 4 thus encoding the transmembrane domain predicted for sTREM2 (*6*). Recently a cleavage site was found on TREM2 that is cleaved by ADAM10, ADAM17, to generate sTREM2 (*20, 21*). The majority of studies of the role of TREM2 in AD pathogenesis have focused on characterizing the functional impact of *TREM2* rare risk variants by using animal and cell models. Some researchers suggest that sTREM2 functions to bind to and sequester TREM2 ligands, such as apolipoproteins, thus preventing them from activating the TREM2 receptor (*47, 48*). Others propose that sTREM2 functions independently of the TREM2 receptor, as the introduction of sTREM2 alters microglia survival and cytokine release in *Trem2^-/-^* cells (*49*).

We present two lines of evidence suggesting that genes within the *MS4A* cluster regulate sTREM2. First, the strength of the signal in *MS4A* gene locus and the consistent association across datasets (*P*_*meta*_ = 4.48×10^−21^; Fig 2). For comparison, the effect size (OR = 3.34; 95% CI = 2.37 - 4.82) of rs1582763 on CSF sTREM2 levels is similar to that of the *APOE* genotype on CSF Aβ_42_ levels (OR = 2.84; 95% CI = 2.30 - 3.50). Second, the *MS4A* association with CSF sTREM2 levels represents a *trans* effect: *TREM2* is located on chromosome 9 and this *MS4A* locus is on chromosome 11. As our group and others have reported, it is not common to find trans-protein quantitative trait loci (pQTLs) with large effect sizes (*35, 50–54*). Most *trans*-pQTLs occur in genes which encode receptors or that have other regulatory influences on the protein studied. For example, variants located on 3p21.31 were significantly associated (rs145617407: *β* = 0.348, *P* = 2.58×10^−10^) with plasma levels of macrophage inflammatory protein 1beta (MIP1b), a protein which is encoded by C-C motif chemokine ligand 4 (*CCL4*) located on 17q12 (*50, 54*). The 3p21.31 locus contains genes which encode five members of the C-C motif chemokine receptor family (CCR1, CCR2, CCR3, CCR5, and CCR9) including a known receptor for CCL4 (CCR5) (*55*). Thus, we hypothesize that one or more genes within the *MS4A* locus are key regulators of sTREM2.

Our study also implicates MS4A4A as a potential therapeutic target for AD. MS4A4A co-localizes with TREM2 in human macrophages and both proteins are upregulated in response to IL-4-mediated stimulation. *TREM2* and *MS4A4A* are highly expressed in human microglia (*37*). While little is known of MS4A4A function in the brain, MS4A4A may be involved in protein trafficking and clathrin-dependent endocytosis (*42*). Our findings that antibody-mediated targeting of MS4A4A was sufficient to reduce sTREM2 levels in human macrophages illustrates a putative relationship between these two proteins. Additional studies are needed to fully understand the mechanism for which MS4A4A affects sTREM2 levels. Compounds that impact MS4A4A levels or block its function could modify AD risk, by regulating brain sTREM2 levels.

## Materials and Methods

### Ethics statement

The Institutional Review Boards of all participating institutions approved the study and research was carried out in accordance with the approved protocols. Written informed consent was obtained from participants or their family members.

### Cohort demographics

Cerebrospinal fluid was obtained from the Alzheimer Disease Neuroimaging Initiative (ADNI) and levels of sTREM2 were measured with enzyme-linked immunosorbent assay (ELISA) by a group of investigators at Washington University in St Louis (WashU) and a group at Ludwig-Maximilians-Universität (LMU) München department of Neurology (Munich, Germany). A total of 207 cognitively normal controls and 606 AD cases were studied in addition to 38 TREM2 mutation carriers (p.D87N, p.H157Y, p.L211P, p.R47H, and p.R62H). TREM2 risk-variant carriers were excluded from the GWAS analyses. Demographic characteristics of the case-control dataset can be seen in Table 1, and Table 2 shows the characteristics of the *TREM2* mutation carriers. Additional datasets were used for replication of the top SNPs, demographic characteristics are shown in Supplementary Table S2. Data was obtained from The Charles F and Joanne Knight Alzheimer’s Disease Research Center (Knight ADRC), GHPH, GHDEM, SPIN, Clinic-IDIBAPS, and The Dominantly Inherited Alzheimer Network (DIAN) (*23–25, 31*).

RNA-seq data were obtained from four independent cohorts as described previously (*36*). The Knight ADRC provided data from 28 brains taken from neuropathologically-confirmed late-onset AD, an additional 10 AD cases with known *TREM2* mutations, and 14 cognitively healthy controls. DIAN provided data from 19 autosomal-dominantly inherited early-onset AD. RNA extraction and sequencing methods were described previously (*36*). RNA-seq data were obtained from the Mayo Clinic Brain Bank via the AMP-AD Knowledge portal (https://www.synapse.org; synapse ID = 5550404; accessed January 2017), the RNA extraction and sequencing methods were described previously (*56*). The AMP-AD portal was also utilized to obtain RNA-seq data from the Mount Sinai Brain Bank (https://www.synapse.org; synapse ID = 3157743; accessed January 2017) which included data from 1,030 samples collected from four brain regions taken from 300 individuals, as described previously (*57*). QC and processing was performing as previously described and additional information is in the supplementary methods (*36*).

### ELISA for CSF sTREM2

CSF samples were obtained and measured separately by investigators at WashU and LMU using two different methods as described below. Measured CSF levels were corrected by plate-specific correction factors by each group and corrected values were used for analyses. Pearson’s correlation was used to compare the corrected sTREM2 values between the two different ELISA methods in overlapping samples (*N* = 980, *r* = 0.834, *P* = 2.2×10^−254^).

The soluble TREM2 ELISA performed at WashU was developed in-house as described previously (*23*) with some changes. Briefly, an anti-human TREM-2 monoclonal antibody (R&D Systems # MAB1828, Clone 263602; 0.5 mg/mL) was used as a capture antibody and coated overnight at 4°C on Maxisorp 96-well plates (Nalge Nunc International, Rochester, NY) in sodium bicarbonate coating buffer (0.015 M Na_2_CO_3_ + 0.035 M NaHCO_3_, pH=9.6). After washing, wells were blocked for 4h at 37°C with PBS 10% fetal bovine serum (FBS). Freshly thawed CSF and recombinant human TREM-2 standard (Sino Biological 11084-H08H-50) were incubated in duplicate overnight at 4°C. For detection, a goat anti-human TREM-2 biotinylated polyclonal antibody (R&D Systems # BAF1828; 0.2 mg/mL) was diluted in assay buffer (PBS 10% FBS at 1:3000) and incubated for 1.25h at room temperature (RT) on orbital shaker. After washing, wells were incubated with horseradish-peroxidase labelled streptavidin (BD Biosciences, San Jose, CA; diluted 1:3000) for 1h at RT with orbital shaking. HRP visualization was performed with 3,3’,5,5’ tetramethylbenzidine (Sigma-Aldrich, St. Louis, MO) added to each well for 10min at RT in the dark. Color development was stopped by adding equal volume of 2.5 N H_2_SO_4_. Optical density of each well was determined at 450 nm. Washes between the different steps were done four times with PBS 0.05% Tween 20 (Sigma-Aldrich). An internal standard, consisting of a single batch of human CSF positive for sTREM-2 was run in all the assays. Samples were run in duplicate in each assay. Raw values are provided as pg/mL. Interassay variability was calculated from duplicate internal standards across 27 plates (CV = 24.01), intra-assay variability from duplicate measurements was < 9.5%.

CSF sTREM2 measurements LMU were done with an ELISA based on the MSD platform and it is comprehensively described in previous publications (*12, 24, 25*). The assay consists of a Streptavidin-coated 96-well plates (MSD Streptavidin Gold Plates, cat. no. L15SA-1); a biotinylated polyclonal goat IgG anti-human TREM2 antibody (R&D Systems, cat. no. BAF1828; 0.25 μg/mL, 25 μL/well) as capture antibody, which is raised against amino acids 19-174 of human TREM2; a monoclonal mouse IgG anti-human TREM2 antibody (Santa Cruz Biotechnology, B-3, cat. no. sc373828; 1 μg/mL, 50 μL/well) as a detection antibody, which is raised against amino acids 1-160 of human TREM2; and a SULFO-TAG-labeled goat polyclonal anti-mouse IgG secondary antibody (MSD, cat. no. R32AC; 0.5 μg/mL, 25 μL/well). All antibodies were diluted in 1% BSA and 0.05% Tween 20 in PBS (pH=7.4) buffer. Recombinant human TREM2 protein (Holzel Diagnostika, cat. no. 11084-H08H), corresponding to the extracellular domain of human TREM2 (amino acids 19-174) was used as a standard (62.5 to 8000 pg/mL). In brief, Streptavidin-coated 96-well plates were blocked overnight at 4°C in blocking buffer [3% bovine serum albumin (BSA) and 0.05% Tween 20 in PBS (pH 7.4); 300 μL/well]. The plates were next incubated with the capture antibody for 1 hour at room temperature (RT). They were subsequently washed four times with washing buffer (200 μL/well; 0.05% Tween 20 in PBS). Thereafter, the recombinant human TREM2 protein (standard curve), the blanks, and the CSF samples and the internal standards IS (duplicates; dilution factor: 4) were diluted in assay buffer [0.25% BSA and 0.05% Tween 20 in PBS (pH=7.4)] supplemented with protease inhibitors (Sigma; Cat. # P8340) and incubated (50 μL/well) for 2 hours at RT. This dilution was previously selected because it showed the best recovery and linearity performance (*12*). Plates were again washed four times with washing buffer before incubation for 1 hour at RT with detection antibody. After four additional washing steps, plates were incubated with SULFO-tag conjugated secondary antibody for 1 hour in the dark at RT. Last, plates were washed four times with wash buffer followed by two washing steps in PBS. The electrochemical signal was developed by adding 150 μL/well MSD Read buffer T (Cat. # R-92TC) and the light emission measured using the MESO QuickPlex SQ 120. Raw values are provided as pg/mL. All CSF samples were distributed randomly across plates and measured in duplicate. The mean intraplate coefficient of variation (CV) was 3.1%; all duplicate measures had a CV < 15%. All the antibodies and plates are commercially available and, for this experiment, they were from a single lot in order to exclude variability between batches. Four internal standards (IS) were run on each ELISA plate to account for interplate variability. All IS used in this study consisted of pooled CSF samples from diagnostic clinical routine leftovers from LMU. The experiment was performed on four different days between the 26th Nov 2017 and 6th Dec 2017 by operators experienced in running these assays and blinded to the clinical information.

### Genotyping and imputation

Samples were genotyped with the Illumina 610 or Omniexpress chip. Stringent quality control (QC) criteria were applied to each genotyping array separately before combining genotype data. The minimum call rate for single nucleotide polymorphisms (SNPs) and individuals was 98% and autosomal SNPs not in Hardy-Weinberg equilibrium (*P* < 1×10^−6^) were excluded. X-chromosome SNPs were analyzed to verify gender identification. Unanticipated duplicates and cryptic relatedness (Pihat ≥0.25) among samples were tested by pairwise genome-wide estimates of proportion identity-by-descent, and when a pair of identical or related samples was identified, the sample from Knight-ADRC or with a higher number of variants that passed QC was prioritized. EIGENSTRAT (*58*) was used to calculate principal components. *APOE* ε2, ε3, and ε4 isoforms were determined by genotyping rs7412 and rs429358 using Taqman genotyping technology as previously described (*59–61*). The 1000 Genomes Project Phase 3 data (October 2014), SHAPEIT v2.r837 (*62*), and IMPUTE2 v2.3.2 (*63*) were used for phasing and imputation. Individual genotypes imputed with probability < 0.90 were set to missing and imputed genotypes with probability ≥0.90 were analyzed as fully observed. Genotyped and imputed variants with MAF < 0.02 or IMPUTE2 information score < 0.30 were excluded, leaving 7,320,475 variants for analyses.

### Statistical methods

Statistical analyses and data visualization were performed in R v3.4.0 (*64*), PLINK v1.9 (*65*), and LocusZoom v1.3 (*66*). Corrected raw values for CSF sTREM2 were normally distributed, so two-sample t-tests were used to compare CSF levels between males and females and between AD cases and controls. Pearson’s product moment correlation coefficient was used for correlation analyses. Single-variant associations with CSF sTREM2 levels were tested using the additive linear regression model in PLINK v1.9 (*65*). Covariates were age at time of LP, sex, and the first two principal component factors to account for population structure. We run a GWAS with the CSF sTREM2 measures from both WashU and LMU. In both cases a genome-wide signal in the MS4 cluster were found (See Figure 1 and Supplementary Figure S3). The genomic inflation factor was λ < 1.008 for all of the genetic analyses. Statistical significance for single-variant analyses was selected based on the commonly used threshold estimated from Bonferroni correction of the likely number of independent tests in genome-wide analyses (*P* < 5 × 10^−8^). To identify additional independent genetic signals, conditional analyses were conducted by adding the SNP with the smallest p-value as a covariate into the default regression model and testing all remaining regional SNPs for association. The dataset was stratified by the most recently reported case status and the genetic signals from the joint dataset were tested to determine whether the genetic associations were driven by cases or controls. To determine whether the identified genetic association within the joint dataset was sex-specific, the dataset was stratified by sex and each sex was tested for the association. Results from the sex-specific analyses were verified in a subset of age-matched samples (*N* = 367 each for males and females, Table S1). Gene expression was inferred using Salmon v0.7.2 (*67*) transcript expression quantification of the coding transcripts of *Homo sapiens* included in the GENCODE reference genome (GRCh37.75). Gene counts data were transformed to stabilize expression variances along the range of mean values and normalized according to library size using the variance stabilizing transformation (VST) function in DESeq2 (*68*). Correlations between expression of *TREM2* and each member of the *MS4A* gene family, with an average TPM > 1, were tested in each independent RNA-seq cohort.

### Bioinformatics annotation

All variants below the threshold for suggestive significance (*P* < 1×10^−5^) were taken forward for functional annotation using ANNOVAR version 2015-06-17 (*69*) and examined for potential regulatory functions using HaploReg v4.1 (*70*) and RegulomeDB v1.1 (*71*). Search tools from publicly available databases, Genotype-Tissue Expression (GTEx) Analysis V7 (*72*), the Brain eQTL Almanac (Braineac) (*73*), and the Westra Blood eQTL browser (*35*), were utilized to determine if significant SNPs were reported eQTLs. The Encyclopedia of DNA elements (ENCODE, https://www.encodeproject.org/) was queried, filtering for brain tissue in *Homo sapiens*, to examine DNA elements affected by the associated SNPs.

### Human macrophages cultures

Peripheral blood mononuclear cells (PBMCs) were purified from human blood on Ficoll-Paque PLUS density gradient (Amersham Biosciences, Piscataway, NJ). To generate macrophages, PBMCs were cultured in 6-well culture plates (3×10^6^ cell/well) in RPMI-1640 without fetal bovine serum. After 2 hours of culture PBMCs were washed twice with PBS 1X and cultured in RPMI supplemented with 50 ng/ml MCSF for 7 days at 37°C with 5% CO_2_. Supernatants from macrophage cultures were collected at different time points, centrifuged at 21000g for 15 min and filtered through a 0.22 βm filter to remove all cells and membrane debris. Macrophage supernatants were then frozen in aliquots at −80°C until used in the ELISA for human sTREM2.

### In vitro experiments with anti MS4A antibodies

Human macrophages after 7 days in culture were incubated with 10 μg/ml of anti-MS4A4 (add here the cat # and brand of anti MS4A4A Abcam (A); Biolegend (B); or anti-MS4A6 (Invitrogen) antibodies in the presence or not of IL-4 (Peprotech; 15 ng/ml) in culture.

After 48 hours cell culture media were collected, centrifuged at 21000g for 15 min and filtered through a 0.22 βm filter to remove all cells and membrane debris. Macrophage supernatants were then frozen in aliquots at −80°C until used in the ELISA for human sTREM2.

### Immunocytochemistry and cell imaging

Cells were fixed for 15-20 min at room temperature (RT) in 4 % (w/v) PFA, 4 % (w/v) sucrose, 20 mM NaOH and 5 mM MgCl2 in PBS, pH 7.4. For intracellular staining, cells were permeabilized and blocked for 60 min at RT in 5% Horse serum, 0.1% Saponin in PBSand incubated at 4 °C overnight with human anti-TREM2 (R&D, AF1828) and human anti-MS4A4A (Biolegend, clone 5C12) primary antibodies diluted in PBS, 5% Horse serum Samples were analyzed with a Zeiss LSM880 laser scanning confocal microscope (Carl Zeiss Inc, Thornwood, NY) equipped with 40X, 1.4 numerical aperture (NA) and 63X, 1.4 NA Zeiss Plan Apochromat oil objectives. ZEN 2.1 black edition software was used to obtain Z-stacks through the entire height of the cells with confocal Z-slices of 1.4μm (40X) and 1.5μm (63X) and an interval 1μm. The 405nm diode, 488nm Argon, and 543nm HeNe1 (helium neon) lasers were utilized with an optimal pinhole of 1 airy unit to acquire images. Images were finally processed with ImageJ software.

## Supplementary Materials

See supplementary materials

## Acknowledgments

We thank all the participants and their families, as well as the many institutions and their staff that provided support for the studies involved in this collaboration.

## Funding

This work was supported by grants from the National Institutes of Health (R01AG044546, P01AG003991, RF1AG053303, R01AG058501, and U01AG058922), the Alzheimer Association (NIRG-11-200110, BAND-14-338165, and BFG-15-362540), YD is supported by an NIMH institutional training grant (T32MH014877). LP was supported by a grant from the Fondazione Italiana Sclerosi Multipla (FISM 2017/R/20). FF was supported by Fondazione Veronesi fellowship. CC was supported during the course of this study by fellowship from the National Multiple Sclerosis Society (FG 2010-A1/2). BAB is supported by 2018 pilot funding from the Hope Center for Neurological Disorders and the Danforth Foundation Challenge at Washington University. The recruitment and clinical characterization of research participants at Washington University were supported by NIH P50 AG05681, P01 AG03991, and P01 AG026276.KB holds the Torsten Söderberg Professorship in Medicine at the Royal Swedish Academy of Sciences. Supported by grants from the Swedish Alzheimer Foundation (# AF-742881), the Research Council, Sweden (#2017-00915), Hjärnfonden, Sweden (# FO2017- 0243), and LUA/ALF project, Vastra Gotalandsregionen, Sweden (# ALFGBG-715986). HZ is a Wallenberg Academy Fellow and is supported by grants from the Swedish and European Research Councils and the UK Dementia Research Institute at UCL. This work was supported by access to equipment made possible by the Hope Center for Neurological Disorders and the Departments of Neurology and Psychiatry at Washington University School of Medicine.

## ADNI

Data collection and sharing for this project was funded by the Alzheimer’s Disease Neuroimaging Initiative (ADNI) (National Institutes of Health Grant U01 AG024904) and DOD (Department of Defense award number W81XWH-12-2-0012). ADNI is funded by the National Institute on Aging, the National Institute of Biomedical Imaging and Bioengineering, and through generous contributions from the following: AbbVie, Alzheimer’s Association, Alzheimer’s Drug Discovery Foundation, Araclon Biotech, BioClinica Inc, Biogen, Bristol-Myers Squibb Company, CereSpir Inc, Cogstate, Eisai Inc, Elan Pharmaceuticals Inc, Eli Lilly and Company, EuroImmun, F. Hoffmann-La Roche Ltd and its affiliated company Genentech Inc, Fujirebio, GE Healthcare, IXICO Ltd, Janssen Alzheimer Immunotherapy Research & Development LLC, Johnson & Johnson Pharmaceutical Research & Development LLC, Lumosity, Lundbeck, Merck & Co Inc, Meso Scale Diagnostics LLC, NeuroRx Research, Neurotrack Technologies, Novartis Pharmaceuticals Corporation, Pfizer Inc, Piramal Imaging, Servier, Takeda Pharmaceutical Company, and Transition Therapeutics. The Canadian Institutes of Health Research is providing funds to support ADNI clinical sites in Canada. Private sector contributions are facilitated by the Foundation for the National Institutes of Health (www.fnih.org). The grantee organization is the Northern California Institute for Research and Education, and the study is coordinated by the Alzheimer’s Therapeutic Research Institute at the University of Southern California. ADNI data are disseminated by the Laboratory for Neuro Imaging at the University of Southern California.

## DIAN

Data collection and sharing for this project was supported by The Dominantly Inherited Alzheimer’s Network (DIAN, UF1AG032438) funded by the National Institute on Aging (NIA), the German Center for Neurodegenerative Diseases (DZNE), Raul Carrea Institute for Neurological Research (FLENI), Partial support by the Research and Development Grants for Dementia from Japan Agency for Medical Research and Development, AMED, and the Korea Health Technology R&D Project through the Korea Health Industry Development Institute (KHIDI). This manuscript has been reviewed by DIAN Study investigators for scientific content and consistency of data interpretation with previous DIAN Study publications. We acknowledge the altruism of the participants and their families and contributions of the DIAN research and support staff at each of the participating sites for their contributions to this study.

## Mayo RNAseq

Study data were provided by the following sources: The Mayo Clinic Alzheimers Disease Genetic Studies, led by Dr. Nilufer Taner and Dr. Steven G. Younkin, Mayo Clinic, Jacksonville, FL using samples from the Mayo Clinic Study of Aging, the Mayo Clinic Alzheimers Disease Research Center, and the Mayo Clinic Brain Bank. Data collection was supported through funding by NIA grants P50 AG016574, R01 AG032990, U01 AG046139,
R01 AG018023, U01 AG006576, U01 AG006786, R01 AG025711, R01 AG017216, R01 AG003949, NINDS grant R01 NS080820, CurePSP Foundation, and support from Mayo Foundation. Study data includes samples collected through the Sun Health Research Institute Brain and Body Donation Program of Sun City, Arizona. The Brain and Body Donation Program is supported by the National Institute of Neurological Disorders and Stroke (U24 NS072026 National Brain and Tissue Resource for Parkinsons Disease and Related Disorders), the National Institute on Aging (P30 AG19610 Arizona Alzheimers Disease Core Center), the Arizona Department of Health Services (contract 211002, Arizona Alzheimers Research Center), the Arizona Biomedical Research Commission (contracts 4001, 0011, 05-901 and 1001 to the Arizona Parkinson’s Disease Consortium) and the Michael J. Fox Foundation for Parkinsons Research.

## MSBB

These data were generated from postmortem brain tissue collected through the Mount Sinai VA Medical Center Brain Bank and were provided by Dr. Eric Schadt from Mount Sinai School of Medicine.

## Author contributions

YD analyzed the GWAS data, performed bioinformatics analyses, and wrote the manuscript. FF, FC, CC, SH, LP and CMK designed and performed the cell-based studies. ZL, JLD, UB, JB, BB and OH performed the gene-expression analyses in human brain tissue. FGF, LI, MVG and JB performed the QC for the GWAS data. KB, HZ, JS, BN, AL, DA, JC, LR, JLM, EMR, and GK provided CSF TREM2 and genetic material for the replication analyses. MSC, CH, YD, LP and CC measured and QCed the CSF sTREM2 in the ADNI dataset. ADNI provided data. LP, CMK, CC supervised and wrote the project. All authors read and approved the manuscript.

## Competing interests

KB has served as a consultant or at advisory boards for Alzheon, BioArctic, Biogen, Eli Lilly, Fujirebio Europe, IBL International, Merck, Novartis, Pfizer, and Roche Diagnostics, and is a co-founder of Brain Biomarker Solutions in Gothenburg AB, a GU Ventures-based platform company at the University of Gothenburg. HZ has served at scientific advisory boards for Eli Lilly, Roche Diagnostics and Wave, has received travel support from Teva and is a co-founder of Brain Biomarker Solutions in Gothenburg AB, a GU Ventures-based platform company at the University of Gothenburg.

## Data and materials availability

All ADNI data are available through the LONI Image & Data Archive (IDA) and interested scientists may apply for access on the ADNI website (https://ida.loni.usc.edu/collaboration/access/).

